# Identifying functional metabolic shifts in heart failure with the integration of omics data and a cardiomyocyte-specific, genome-scale model

**DOI:** 10.1101/2020.07.20.212274

**Authors:** Bonnie V. Dougherty, Kristopher D. Rawls, Glynis L. Rolling, Kalyan C. Vinnakota, Anders Wallqvist, Jason A. Papin

**Affiliations:** Department of Biomedical Engineering, University of Virginia, Charlottesville, VA, 22908, USA; Department of Medicine, Division of Infectious Diseases and International Health, University of Virginia, Charlottesville, VA, 22908, USA; Department of Defense Biotechnology High Performance Computing Software Applications Institute, Telemedicine and Advanced Technology Research Center, U.S. Army Medical Research and Development Command, Fort Detrick, Maryland, 21702 USA; The Henry M. Jackson Foundation for the Advancement of Military Medicine, Inc., Bethesda, Maryland, 20817, USA; Department of Biochemistry & Molecular Genetics, University of Virginia, Charlottesville, VA, 22908, USA

**Keywords:** metabolic network, cardiac metabolism, heart failure

## Abstract

The heart is a metabolic omnivore, known to consume many different carbon substrates in order to maintain function. In diseased states, the heart’s metabolism can shift between different carbon substrates; however, there is some disagreement in the field as to the metabolic shifts seen in end-stage heart failure and whether all heart failure converges to a common metabolic phenotype. Here, we present a new, validated cardiomyocyte-specific GEnome-scale metabolic Network REconstruction (GENRE), *iCardio*, and use the model to identify common shifts in metabolic functions across heart failure omics datasets. We demonstrate the utility of *iCardio* in interpreting heart failure gene expression data by identifying Tasks Inferred from Differential Expression (TIDEs) which represent metabolic functions associated with changes in gene expression. We identify decreased NO and Neu5Ac synthesis as common metabolic markers of heart failure across datasets. Further, we highlight the differences in metabolic functions seen across studies, further highlighting the complexity of heart failure. The methods presented for constructing a tissue-specific model and identifying TIDEs can be extended to multiple tissue and diseases of interest.

## Introduction

In order for the heart to maintain its function, the main contractile unit (cardiomyocyte) uses oxygen and various nutrients through metabolic pathways to maintain contractile proteins, synthesize necessary lipid species, and produce ATP as a fuel for muscle contraction. Metabolic changes, such as changes in substrate utilization, have been noted in many diseased states, such as left ventricular hypertrophy (Kundu et al., 2015) and cardiotoxicity (Bauckneht et al., 2017; Borde et al., 2012). In some cases, changes in substrate utilization occur before functional and/or structural changes to the heart, suggesting that metabolism plays a key role in the downstream development of disease or could be a target to prevent disease (Kundu et al., 2015; Li et al., 2019). However, it is not understood if changes in cardiomyocyte metabolism are the result of, a contributor to, or the cause of disease. Therefore, there is a need for a comprehensive, descriptive model of the metabolic function of the heart to interrogate the relationships between substrates and function.

A common tool to interrogate relationships between metabolic substrates and metabolic function is a GEnome-scale metabolic Network REconstruction (GENRE). GENREs are mathematical representations of metabolism that use the enzymes encoded in an organism’s genome to define the biochemical metabolic reactions and associated metabolites that comprise that organism’s metabolism. Each metabolic reaction is associated with a gene-protein-reaction (GPR) rule relating genes to the proteins they encode and proteins to the reactions they catalyze. Human GENREs account for the function of the biochemical reactions that humans catalyze according to annotation of the human genome.

However, not all genes are expressed in every tissue, necessitating the construction of tissue-specific models of metabolism. GENREs of human metabolism (Blais et al., 2017; Brunk et al., 2018; Duarte et al., 2007; Ma et al., 2007; Mardinoglu et al.; Swainston et al., 2016; Thiele et al., 2013) have been used to generate tissue-, disease-, or cell-specific models for various analyses such as predicting drug targets (Agren et al., 2014; Folger et al., 2011; Rawls et al., 2020), identifying biomarkers of disease (Zhang et al., 2013; Zhao and Huang, 2011), and understanding drug toxicity or side effects (Blais et al., 2017; Shaked et al., 2016; Zielinski et al., 2015). Tissue-specific models are built by integrating tissue-specific omics data, usually transcriptomic or proteomic data, with a human GENRE to obtain a tissue- or cell-type specific model using various integration algorithms (some of which are summarized in (Blazier and Papin, 2012; Robaina Estévez and Nikoloski, 2014)). To date, there are two existing cardiomyocyte models (Karlstädt et al., 2012; Zhao and Huang, 2011), both of which were built from Recon1, the first human GENRE (Duarte et al., 2007). These models were used to examine the relationship between substrate utilization and efficiency of the heart (Karlstädt et al., 2012) and to predict epistatic interactions in the heart and biomarkers of heart disease (Zhao and Huang, 2011). However, human models have been expanded since Recon1 to more comprehensively describe human metabolism. Our recently published human GENRE, *iHsa* (Blais et al., 2017), is more comprehensive than Recon1 (8336 reactions vs. 3311 reactions) and therefore offers the potential to generate a more comprehensive cardiomyocyte-specific model. A new cardiomyocyte-specific model can be used as a tool to interpret large data sets, such as transcriptomic data, to provide functional insight into how metabolism might change in a diseased state.

Here, we present a new, validated, cardiomyocyte-specific metabolic model, *iCardio*, which was built using *iHsa* (Blais et al., 2017) with data from the Human Protein Atlas (HPA) (v18.proteinstlas.org; Uhlén et al., 2015). The draft model was curated using metabolic tasks, which are mathematical descriptions of metabolic functions that a model should be able to perform. The metabolic tasks represent both previously published tasks (Blais et al., 2017) as well as newly developed, cardiomyocyte-relevant tasks. We validated the model qualitatively by the number of reactions covered by the metabolic tasks and quantitative ATP predictions for common carbon sources.

Finally, we demonstrate the utility of metabolic models in identifying functional changes in metabolism using a new method that identifies metabolic tasks associated with significant changes in gene expression, called <u>T</u>asks <u>I</u>nferred from <u>D</u>ifferential <u>E</u>xpression (TIDEs). TIDEs are a unique type of gene set enrichment analysis (GSEA) that take into account both the stoichiometric balance of reactions necessary to achieve metabolic functions by including transporter reactions between metabolic compartments as well as the complex relationship between genes, proteins, and the reactions they catalyze. Therefore, TIDEs offer biological insight into metabolic functions affected in a disease state that are not readily apparent from gene expression data or gene set enrichment analyses alone. We use heart failure as a case study because of the critical role that metabolism plays in the progression and diagnosis of disease.

## Results

### Building and validating iCardio using metabolic tasks

The CORDA algorithm was used to build the draft *iCardio* model from *iHsa*; CORDA removed 4203 reactions from *iHsa* to build a draft *iCardio* model that had a cardiomyocyte-specific task accuracy of 89% (Figure 1B-C, Supplemental File 2). The draft model was then curated using metabolic tasks to obtain the final *iCardio* model (Figure 1). To achieve 100% task accuracy, 79 reactions were added to and 12 removed from the draft *iCardio* model (Figure 1A) based on literature and manual curation (Supplemental File 2). The 79 reactions that were added to *iCardio* are reactions that were removed from *iHsa* to build the *iCardio* draft model were added back to the *iCardio* model to ensure functionality. We can map the protein data from HPA using the associated GPR rules. Of the 79 reactions added back to *iCardio*, 73% (58) are associated with either high (1), medium (8), or low (16) protein evidence, no GPR rule (28), or no data (5); the remaining 21 reactions were associated with no protein evidence. As an example, the synthesis of nitric oxide was a metabolic task that originally failed but was curated to pass in the final *iCardio* model. For this task to pass, three reactions were added back to the model, two reaction that were associated with proteins that were not detected and one that had no GPR rule (Supplemental File 2). All three isoforms of nitric oxide synthase (NOS), which catalyze the two reactions, were not detected in v18 of the HPA tissue data but two of the NOS isoforms, NOS1 and NOS2, are associated with low expression in v19, while NOS3 has significantly higher transcript counts in heart muscle in the RNA consensus tissue HPA data (Uhlén et al., 2015).

**Figure 1.**
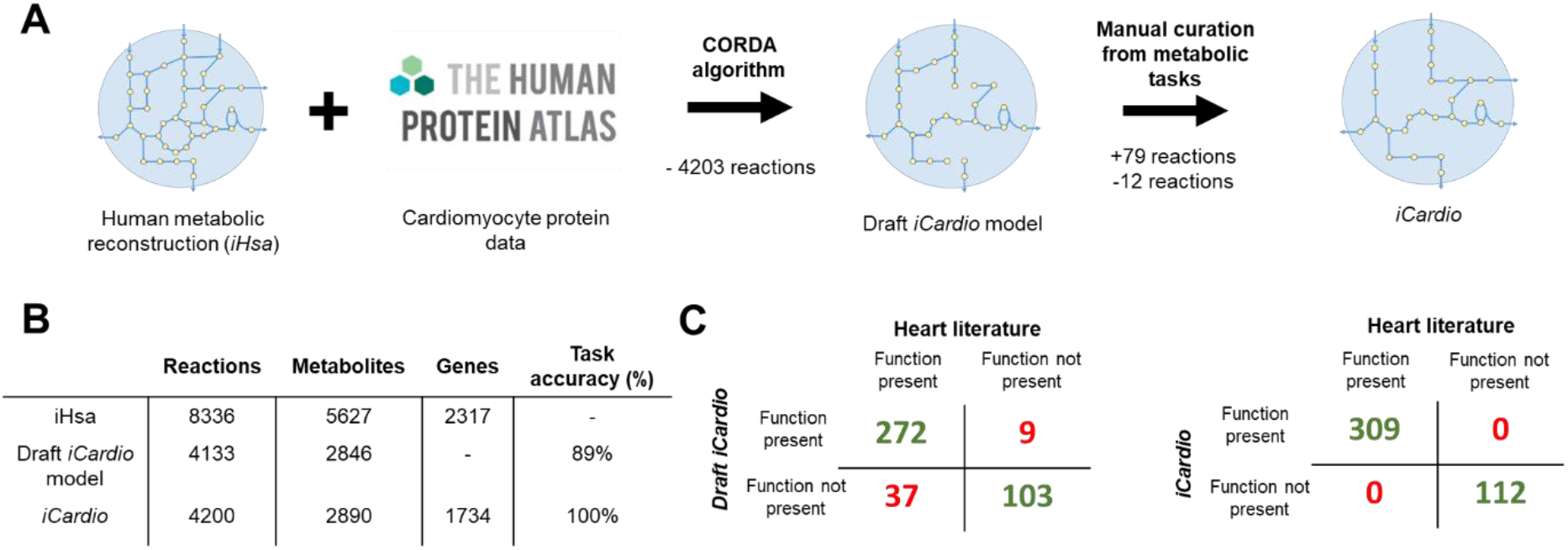
Building a cardiomyocyte metabolic model (iCardio) by integrating protein data and curating with metabolic tasks. (A) The draft iCardio model was built by integrating protein data from the human protein atlas (HPA) with a human metabolic reconstruction (iHsa) using the CORDA algorithm. The draft iCardio model was curated using pre-defined metabolic tasks, resulting in the final model, iCardio. (B) The CORDA algorithm removed 4203 from iHsa to produce a draft iCardio model that had a cardiomyocyte-specific task accuracy of 89%. (C) The superset of 421 metabolic tasks represented metabolic tasks from a previous publication (Blais et al, 2017) with additional metabolic tasks that were curated from the literature based on cardiac metabolism. These tasks were used to test if the model could perform functions known to be present in the heart while removing metabolic functions not present in the heart. After curation, the final iCardio model has a 100% cardiomyocyte-specific task accuracy.

Of note, after curation, the final *iCardio* model contains approximately 50% of the reactions that are present in the *iHsa* network reconstruction. Using the complex GPR rules that are associated with each reaction, we can also identify the number of genes that are associated with a reaction in *iHsa* but are not associated with a reaction in the *iCardio* model; only 583 genes are associated with a reaction in *iHsa* but not included in *iCardio*, highlighting both the promiscuity of enzymes and the complex relationship between genes and metabolic functions (Figure 1B).

Since metabolic tasks have been used as a metric for building *iCardio*, the number of reactions covered by this set of metabolic tasks provides a qualitative validation of the model. As has been done with other models (Richelle et al., 2019), pFBA was used to determine the reactions that each passing metabolic task utilized in *iCardio*. The 216 previously published cardiomyocyte-relevant tasks (Blais et al., 2017) covered 38% (1593/4200) of reactions in *iCardio* and the 93 passing new cardiomyocyte-relevant tasks covered 21% (874/4200) of reactions in *iCardio* (Figure 2A). There is overlap between these two sets of reactions; in total, the two sets of tasks covered 41% (1714/4200) of reactions in *iCardio*. It is important to note that although the tasks may cover the same reactions, they cover different combinations of reactions and pathways for each task. The two sets of tasks together may cover 1,700 reactions but in total over 20,000 reactions are used in different combinations to complete the tasks, indicating that a number of reactions are repeated between tasks. This result is to be expected, especially for reactions that involve central carbon metabolism and ATP production, such as ATP synthase. The maximum number of reactions covered by a given task was 788 reactions (1 task), the metabolic task that describes the de novo synthesis of lipids from glucose and essential fatty acids, and the minimum number of reactions covered by a task was one reaction (24 tasks), metabolic tasks that describe transport. Overall, the reaction coverage of tasks demonstrates a qualitative validation of *iCardio*.

**Figure 2.**
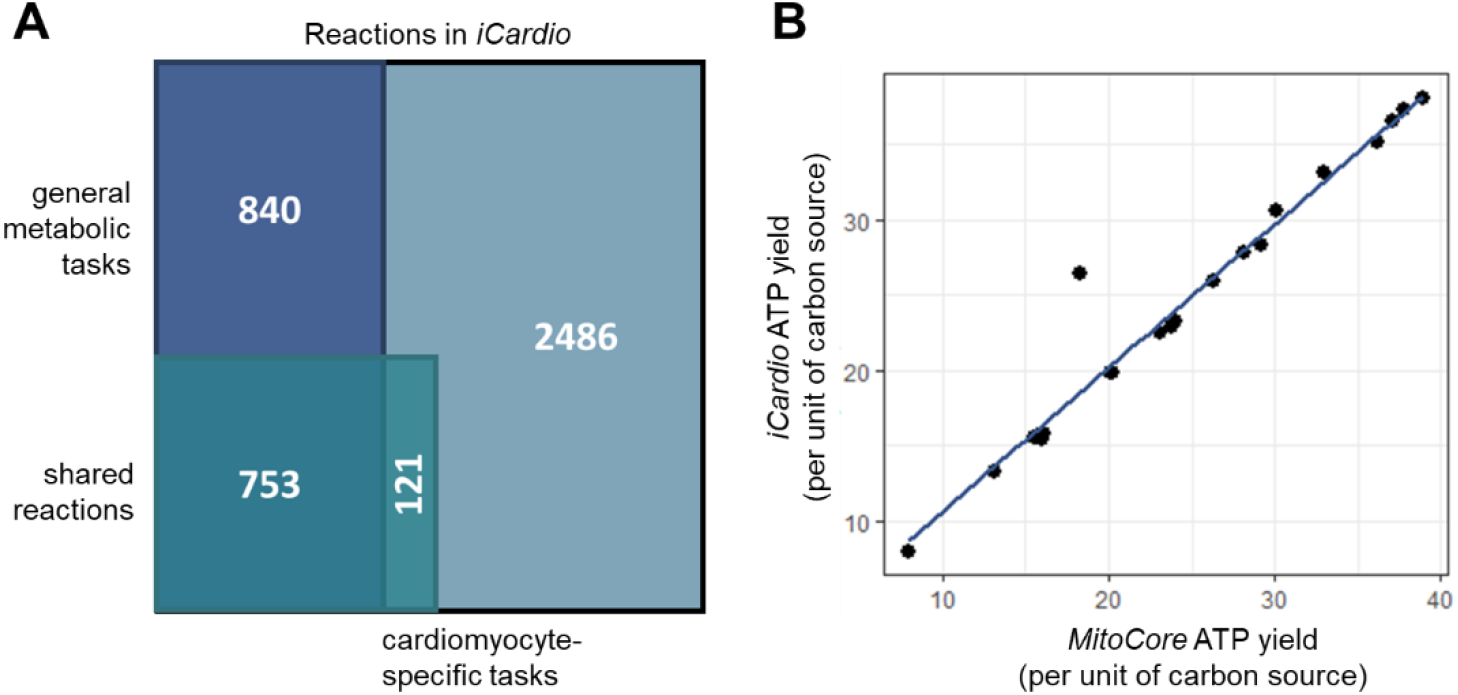
Validating iCardio (A) descriptively using the number of reactions covered my metabolic tasks and (B) quantitatively using ATP yields for common carbon substrates. (A) A descriptive validation of iCardio using the number of reactions covered using the different metabolic tasks. Together, the tasks account for almost half of the reactions in iCardio. The remaining reactions represent areas for future improvement of metabolic tasks. (B) A qualitative validation of iCardio using ATP yields for a variety of common carbon sources. The ATP yields for iCardio (y-axis) are compared to another recently published, but smaller, metabolic model, MitoCore (Smith et al., 2017), which contains 324 reactions. The agreement between the models demonstrates the lack of energy generating cycles and infinite loops in iCardio.

To provide a quantitative validation of *iCardio*, ATP yields were predicted for a number of carbon substrates and amino acids and compared to another metabolic model, *MitoCore* (Smith et al., 2017). The *MitoCore* model was chosen for comparison because of its focus on cardiomyocyte mitochondrial metabolism. For almost all carbon sources (other than methionine), *iCardio* predicted ATP yields within 10% of the values calculated with *MitoCore* (Smith et al., 2017) (Figure 2B). For methionine, the ATP prediction from *iCardio* matches the methionine prediction from Recon 2.2 (Swainston et al., 2016). It is important to note the difference in scope between the two models: *MitoCore* contains 342 reactions whereas *iCardio* contains 4200 reactions while still maintaining accurate ATP yields. This result highlights that, even with the increased size of *iCardio*, there are not infeasible energy-generating loops, which would artificially inflate ATP yield predictions and influence the reactions necessary for different metabolic tasks. Together, the qualitative and quantitative validation of *iCardio* demonstrate the ability of the model to accurately and more comprehensively represent cardiomyocyte metabolism.

### *Identifying <u>T</u>asks <u>I</u>nferred from <u>D</u>ifferential <u>E</u>xpression (TIDEs) using* iCardio *for heart failure gene expression data*

Metabolic models provide an alternative approach for interpreting changes in gene expression to yield insight into metabolic shifts that may be contributing to a diseased state. Here, we use *iCardio* to identify metabolic tasks that were significantly associated with differentially expressed genes, called <u>T</u>asks <u>I</u>nferred from <u>D</u>ifferential <u>E</u>xpression (TIDEs), which represent metabolic functions that are shifted in an experimental versus control state (Figure 3). Heart failure is a complex disease, both in etiology and presentation, but heart failure is associated with a shift in metabolism (Wende et al., 2017). Several different metabolic shifts have been noted in heart failure, such as decreased fatty acid utilization (Lopaschuk, 2017; Wende et al., 2017), increased ketone body utilization (Janardhan et al., 2011), and decreased utilization of branched chain amino acids (Sun et al., 2016). However, there are still disagreements in the field. For example, one study noted an increase in fatty acid uptake in heart failure (Taylor et al., 2001), while other studies reveal decreased fatty acid utilization (Doenst et al., 2013). *iCardio* provides an opportunity to contextualize gene expression data from end-stage heart failure patients to identify changes in metabolic functions, such as those listed above, that are associated with significant changes in gene expression.

**Figure 3.**
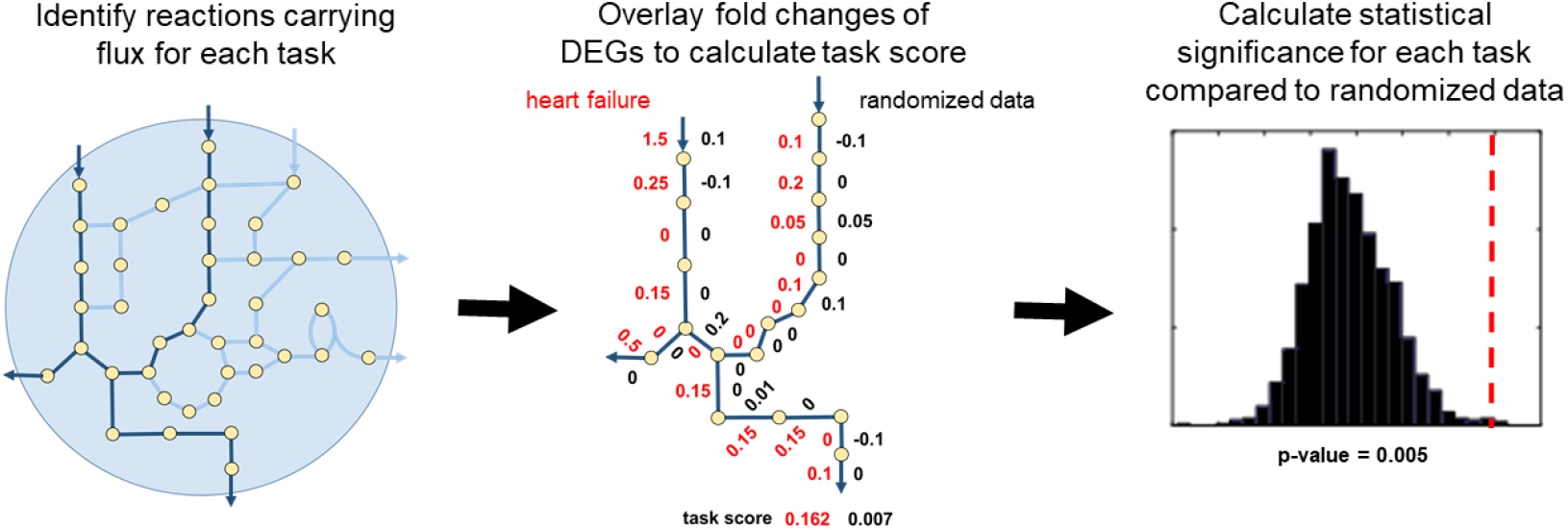
Identifying functional metabolic changes from gene expression data using metabolic tasks. Metabolic tasks quantitatively describe metabolic functions that a tissue or organism is known to catalyze. iCardio was used to identify the reactions that are utilized in order to perform each of the previously defined metabolic tasks. Next, each reaction is assigned a reaction weight based on gene expression fold changes. A task score is calculated based on the weights for each reaction and represents an average gene expression value over reactions utilized in that metabolic task. In order to determine the statistical significance of this task score, the original gene expression data is shuffled over all genes with data and task scores are recalculated to give a distribution of task scores. A p-value is assigned based on where the original task score falls in the distribution of randomized data.

To do this, we analyzed gene expression data from patients undergoing heart transplants for either advanced ischemic or idiopathic heart failure compared to healthy hearts. We integrated these differentially expressed genes (DEGs) and their associated log fold changes with *iCardio* to determine TIDEs, representing metabolic functions associated with a significant change in gene expression. When compared to healthy hearts, the ischemic hearts from GSE5406 had 2678 DEGs; 392 of these DEGs were present in *iCardio* (Supplemental Table 3). After integrating these DEGs using *iCardio* and the TIDEs pipeline, 94 of the 307 metabolic tasks were designated as TIDEs (Figure 4), representing metabolic functions associated with significant changes in gene expression in these heart failure conditions. In the randomized data used to determine the statistical significance of each task, only 5 out of the 1000 random iterations had more than 94 tasks identified as significantly changed, indicating that the identified TIDEs cannot be attributed to random changes in gene expression, but rather distinct and coordinated shifts in gene expression resulting in changes in specific metabolic functions. Across the 307 metabolic tasks, there are varied distributions in the underlying randomized task scores (Figure 4), representing the complex relationships between gene expression and metabolic function that are captured using *iCardio* and the TIDEs reaction-centric approach.

**Figure 4.**
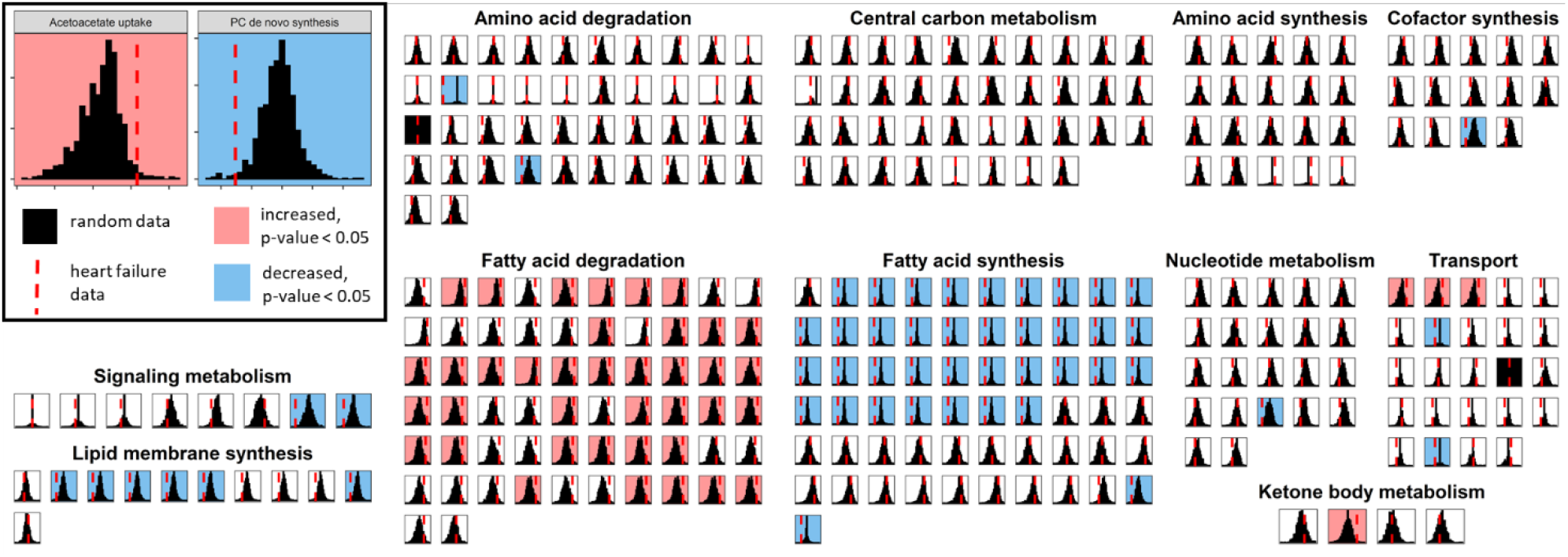
Distributions for model-predicted task scores from gene expression data for ischemic heart failure for each of the 307 tested tasks. Gene expression data for ischemic heart failure vs. healthy hearts from GSE5406 was integrated with iCardio to identify shifts in metabolic functions. The red dashed lines indicate the task score for the heart failure dataset and the black histograms are the distribution of randomized task scores. A red line to the right indicates a metabolic function associated with increased gene expression whereas a red line to the left indicates a metabolic function associated with decreased gene expression. The color of the background indicates if the metabolic function had a statistically significant (p-value < 0.05) increased (red) or decreased (blue) task score.

Some general trends appear from the TIDEs identified from the ischemic heart failure data (Figure 4). First, metabolic tasks related to fatty acid degradation (41) were increased in the heart failure dataset compared to randomized data while metabolic tasks related to fatty acid synthesis (38) were decreased. Within tasks related to fatty acid synthesis, significant decreases were observed for saturated fatty acids but not for unsaturated fatty acids. Finally, metabolic tasks related to lipid membrane synthesis were decreased (6). Taken together, these metabolic shifts seem to support the common hypothesis of heart failure as a state of increased demand for ATP (Ingwall, 2004; Neubauer, 2007), seen through an increased degradation of fatty acids, while limiting other uses of ATP and carbon sources for other functions, such as synthesis of fatty acids and components of the lipid membrane. However, other previously reported metabolic signatures of heart failure, such as increased ketone body degradation (Janardhan et al., 2011), and decreased breakdown of branched chain amino acids (Sun et al., 2016), were not identified using this dataset and approach.

Next, we expanded the TIDE analysis to the two remaining studies (Supplemental Table 3), including both idiopathic and ischemic heart failure. Here, there is no consistent TIDE across all 6 datasets (Figure 5A). However, there were a few frequently observed changes in TIDEs across the datasets. First, nitric oxide synthesis from arginine is decreased in 5 of the 6 datasets (excluding the ischemic heart failure data from GSE1869). Synthesis of long-chain, unsaturated fatty acids was decreased in 4 of the 6 datasets (excluding both ischemic and idiopathic heart failure data from GSE1869). De novo synthesis of Neu5Ac was decreased in 4 of the 6 datasets (excluding the ischemic and idiopathic heart failure data from GSE57345). The breakdown of valine to succinyl-CoA was increased in 2 of the 6 datasets (the idiopathic heart failure data from GSE1869 and GSE5406) and the breakdown of threonine to a Krebs cycle intermediate was increased in 2 of the 6 datasets (both ischemic and idiopathic heart failure in GSE57345). Finally, acetoacetate uptake, B-hydroxybutyrate uptake, and breakdown of B-hydroxybutyrate to acetyl-CoA, representing ketone body metabolism, were increased in 2 of the 6 datasets (both ischemic and idiopathic heart failure data from GSE5406). Across the different datasets, the previously reported metabolic signatures of heart failure appear. However, they tend to appear within a study rather than across all studies or types of heart failure. Finally, rather than clustering by type of heart failure (ischemic vs idiopathic), the TIDEs results cluster by the study.

**Figure 5.**
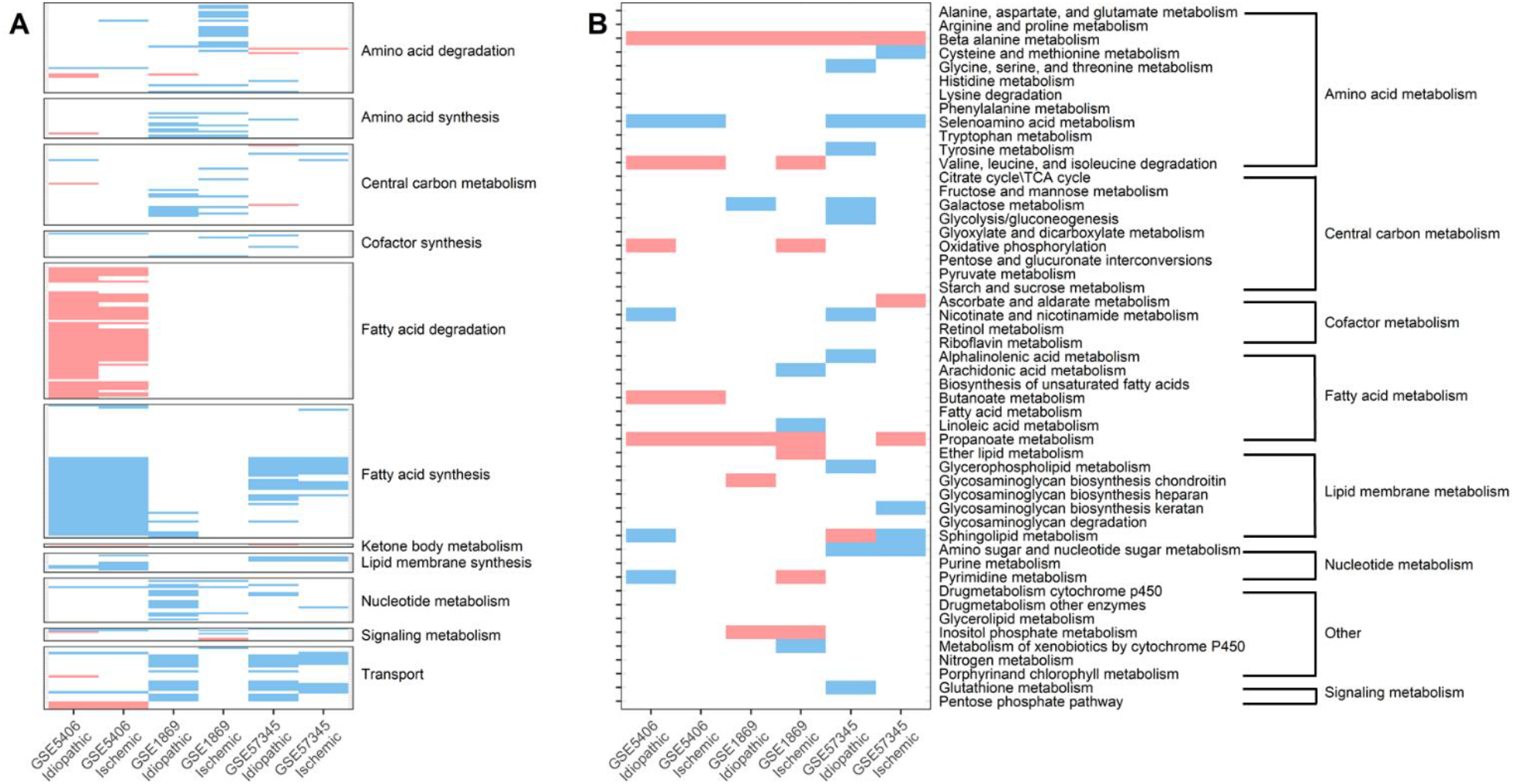
TIDEs analysis identifies functional metabolic changes across different datasets (A) compared to results from a traditional GSEA analysis (B). (A) Three datasets that contained samples for ischemic, idiopathic, and healthy hearts were downloaded, analyzed, and integrated with iCardio using the TIDEs pipeline to identify shifts in metabolic functions. For each of the three datasets, both ischemic and idiopathic heart failure are shown. TIDEs cluster by study rather than type of heart failure. (B) The same three datasets were used with a traditional GSEA analysis with gene sets defined by KEGG pathways. The KEGG metabolic pathways are shown here and grouped similar to the TIDE groupings shown in (A).

### Comparison of TIDEs with GSEA by KEGG pathway

Next, we wanted to compare the TIDEs analysis to a common gene-centric approach, gene set enrichment analysis (GSEA), using KEGG pathways to define the gene sets. Rather than taking a reaction-centric approach, GSEA takes a gene-centric approach and defines a group of genes for specific functions (Subramanian et al., 2005). While KEGG includes many other pathways other than metabolism (Supplemental Figure 1), we have chosen to focus on a comparison of metabolic pathways. Using GSEA with the data from the studies cited above (Figure 5B), the most commonly changed metabolic pathways across the datasets were increased beta alanine metabolism in all datasets, increased propanoate metabolism in 5 of the 6 datasets, decreased selenoamino acid metabolism in 4 of the 6 datasets and increased valine, isoleucine, and leucine degradation in 3 of the 6 datasets. For the commonly cited metabolic shifts in heart failure, the GSEA analysis shows no change in fatty acid metabolism, one dataset with a decrease in genes associated with glycolysis, two studies that show increased oxidative phosphorylation, and 3 datasets with increased valine, isoleucine, and leucine metabolism. For comparison with the metabolic tasks, the breakdown of propanoate and synthesis of beta-alanine were included as metabolic tasks task but showed no change across datasets. Selenoamino acids are included in the model but are not included as a metabolic task. Valine, isoleucine, and leucine metabolism cover six metabolic tasks in the model covering uptake and breakdown of each amino acid. Valine, isoleucine, and leucine were associated with decreased uptake in idiopathic heart failure for GSE1869 and GSE57345. Breakdown of valine to succinyl-CoA was increased in idiopathic heart failure for GSE1869 and GSE5406 while the breakdown of isoleucine to acetyl-CoA and succinyl-CoA was increased in idiopathic heart failure for GSE5406.

The difference in identified changes in metabolism highlight the differences between a reaction-centric approach, such as TIDEs, versus a gene-centric approach. GSEA KEGG metabolic pathways can cover more than one metabolic function, such as fatty acid metabolism, while the TIDEs approach allows for a distinction between the synthesis and degradation of different fatty acids. We can use *iCardio* for a similar approach by using the metabolic subsystems assigned to each reaction to identify a set of reactions. Using TIDEs, we can determine if the reactions in a given subsystem are associated with increased or decreased gene expression (Supplemental Figure 2). While this approach also reveals some interesting trends, such as a decrease in expression for reactions in the arachidonic acid metabolic subsystem, it fails to highlight changes in fatty acid metabolism or other smaller pathways, such as nitric oxide synthesis or ketone body degradation.

Second, a GSEA approach includes every gene that can be present in the pathway even if the function is redundant, such as with isozymes. For example, take the metabolic task of synthesis of nitric oxide from arginine (Figure 6). Using the TIDEs approach with iCardio, this metabolic task covers 5 reactions, 4 of which are associated with GPR rules and therefore included in downstream analysis (Figure 6A). Two of these reactions can be catalyzed by one of three enzymes, NOS1, NOS2, or NOS3, which are known to have tissue-specific expression. The TIDEs reaction-centric approach uses the log-fold change of one gene to determine the reaction weight, which could differ between studies and across tissues. In this study, the two reaction weights for the transporters responsible for the import of arginine to and the export of citrulline from the cytosol are determined by different genes between the idiopathic heart failure data in GSE5406 and GSE57345 (Figure 6C). However, the metabolic task of nitric oxide production from arginine was associated with decreased gene expression in both datasets. A similar GSEA analysis would have included all 6 genes rather than the 2-3 used in the TIDEs analysis.

**Figure 6.**
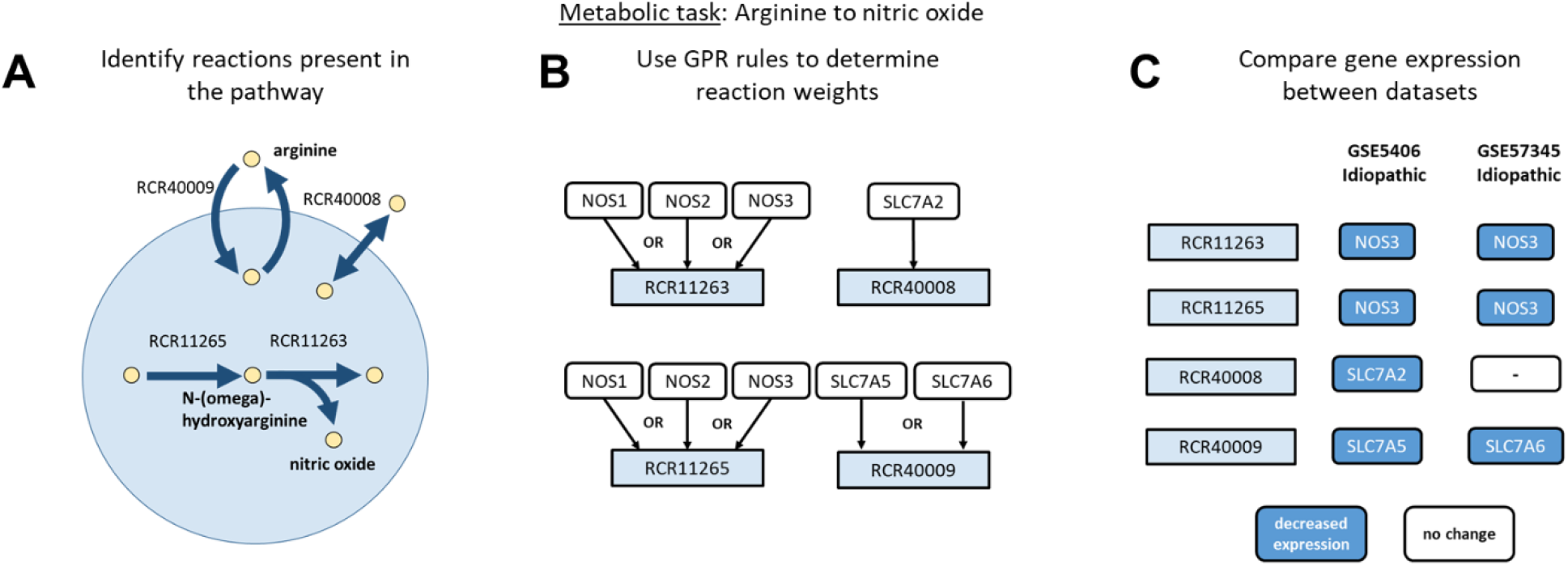
TIDEs analysis identifies differences in datasets for genes which catalyze the conversion of arginine to nitric oxide. (A) For the metabolic task of arginine to nitric oxide, the iCardio model uses four reactions, which cover the transport of arginine and citrulline into the cell and the conversion of arginine to the intermediate N-(omega)-hydroxyarginine and finally conversion to nitric oxide. (B) The model uses the GPR rules associated with each of these reactions to determine the reaction weight based off gene expression data, thereby assigning the expression of one gene to govern the reaction. (C) The gene whose gene expression was used in the analysis can therefore differ between different datasets while still producing the same result, as seen here with both datasets still showing a statistically significant decrease the metabolic function of conversion of arginine to nitric oxide.

## Discussion

Here, we present a new, validated model of cardiomyocyte metabolism, *iCardio*, with a method for analyzing changes in metabolic functions based on gene expression data. We present a new approach for building tissue-specific models using tissue-specific protein data from the HPA integrated with a general human reconstruction, *iHsa*, followed by manual curation with metabolic tasks to ensure general metabolic functionality. Using a task-driven approach for model curation demonstrated that the CORDA method produced a draft model that was more accurate with respect to performance of cardiomyocyte-specific metabolic tasks than other integration methods (Supplemental Table 2). It is interesting to note that the CORDA algorithm produced one of the smaller draft models while still maintaining the highest task accuracy. As noted, although the CORDA algorithm produced one of the smaller draft models and resulted in the final *iCardio* model that contained 50% of the reactions in *iHsa*, 75% of the genes in *iHsa* are still represented in the *iCardio* model. This result highlights the ability of the complex GPR rules to capture tissue-specific function. For example, the conversion of arginine to nitric oxide (NO) is an established mechanism for intracellular and extracellular signaling and the production of NO is mediated through tissue-specific expression of different NOS isoforms (Figure 6). *iCardio* captures the complex relationship between gene expression and function through the complex GPR rules even though all of the isoforms were not detected in the original HPA dataset.

The metabolic tasks used for benchmarking the integration algorithms and further manual curation cover 41% of the network while also identifying areas for future development of both general and tissue-specific metabolic tasks. Here, metabolic tasks were formulated agnostic to the gene expression data that was integrated with the resulting model. Additional tasks could serve as hypotheses for changes in metabolic functions of the heart that could then be tested with specific gene expression data sets. Finally, although not every reaction is covered by a metabolic task, these reactions represent metabolic functions with either evidence of protein expression in the HPA or pathway connectivity from the CORDA integration algorithm. These reactions and associated pathways further support the use of metabolic network reconstructions for generating hypotheses for important, tissue-specific metabolic functions.

The pFBA approach assumes that the pathway for each metabolic task remains the same, independent of the data. This reaction-centric TIDE approach for determining significantly changed metabolic tasks emphasizes that (a) metabolic functions require multiple complex, stoichiometrically balanced reactions in order to proceed, and (b) some genes may disproportionately influence the completion of metabolic functions. For example, two metabolic tasks cover the synthesis of nitric oxide from arginine (Supplemental File 1). The first task comprehensively covers the entire pathway, providing only arginine and other ions extracellularly as inputs while requiring the production of nitric oxide and the release of only metabolites that have transport reactions from the cytosol to the extracellular space. The second task only accounts for the central reactions for nitric oxide production, providing arginine and NADPH and require the production of nitric oxide. While both tasks cover the synthesis of nitric oxide from arginine, the first metabolic task uses 66 reactions and covers the entire pathway, including transport of metabolites between compartments, which are necessary to maintain the stoichiometric balance of the pathway. However, because the second metabolic task covers mainly reactions in the cytosol, it requires only 5 reactions. While the first task was present in the original *iHsa* task, the second task was added because of the importance of the synthesis of nitric oxide in the heart and therefore covers the core reactions for nitric oxide synthesis. Together, these two tasks highlight the potential for both breadth and specificity in identifying shifts in metabolic functions.

Second, reactions are associated with complex GPR rules, allowing for multiple genes representing different isozymes to catalyze the same reaction. By accounting for these complex relationships in the calculation of task scores, the expression values of individual genes can affect multiple functions. For example, fatty acid synthase, which catalyzes multiple reactions, can disproportionately influence a final task score. The associated fold change for fatty acid synthase may determine the reaction weight for multiple reactions in a metabolic task. In contrast, previous gene-centric approaches would have evenly weighted all genes in a specific pathway. The TIDEs analysis represents a reaction-centric approach that focuses on metabolic functions that can provide insight into broad metabolic changes that may not be immediately apparent using gene-centric approaches.

The presented TIDEs pipeline offers an alternative, reaction-centric approach to interpret complex changes in gene expression data to identify non-obvious changes in metabolic functions. In the case of heart failure, the TIDEs pipeline was able to identify some of the common shifts noted in heart failure in at least one of the three studies, such as increased fatty acid utilization, increased utilization of branched chain amino acids, and increased utilization of ketone bodies. The TIDEs analysis also revealed two changes that were common across more than one dataset: decreased synthesis of nitric oxide from arginine and decreased de novo synthesis of Neu5Ac. With an evaluation of all six data sets, decreased expression of NOS3, also referred to as eNOS, was driving the weight of two of the reactions in the metabolic task (Figure 6). Previous work has highlighted the important role of NO in cardiac function (Li et al., 2020; Massion et al., 2003) and more recent work has suggested a role for increasing NO synthesis for the increasing efficacy of beta-blockers (Hayashi et al., 2018). Neu5Ac is the most common sialic acid associated with multiple functional roles in the body. Studies have shown that increased Neu5Ac is associated with the development of atherosclerosis and synthesis of Neu5Ac has been suggested as a therapeutic target (Zhang et al., 2019). Using TIDEs, we can identify genes that are driving the change in the metabolic task, both genes that are shared across and are different between studies which can serve as a starting point for future work. However, more work is needed to identify the role that decreased synthesis of Neu5Ac may play in end-stage heart failure. Both NO and Neu5Ac synthesis represent potential biomarkers of and targets for treating heart failure.

Although there were some common trends across datasets, no TIDEs could discriminate between ischemic and idiopathic heart failure across studies. This characteristic was also true for the KEGG GSEA results. Together, this result highlights the complexity of heart failure, both in etiology and presentation, suggesting that classifications such as ischemic and idiopathic may be insufficient to capture distinct metabolic changes. In addition, the datasets cluster within each study for both the TIDEs and GSEA KEGG metabolic pathway analysis, suggesting, again, that there is a large amount of heterogeneity in changes in gene expression for heart failure. However, it is important to note that changes in gene expression are not the only drivers of changes in metabolic function. Other studies have noted the role of changing metabolic milieu in the blood as a driver of changes in the metabolic functions of the heart during heart failure (Neubauer, 2007). Future work can integrate clinical measures, such as LVEF or FDG-PET glucose uptake measures, which could help to separate clusters of patients and more clearly identify the influence of metabolism in the progression of heart failure.

Here, we provide a validated approach for constructing a tissue-specific metabolic model and demonstrate the utility of metabolic models to interpret changes in metabolic functions based on gene expression data. The model-building process can be extended to other cell- or tissue-type specific models. The metabolic tasks provide a two-fold role for model validation and concrete metabolic functions to identify metabolic shifts in gene expression data. TIDEs represent a new, reaction-centric approach to identifying changes in metabolic functions and testing new hypotheses for changes in gene expression for metabolic functions. These new hypotheses can be formulated as metabolic tasks based on the current literature, based on reactions present in a metabolic model but not covered in the current list of metabolic tasks, or results from other gene-centric approaches, such as GSEA. The method is not limited to use with *iCardio* but can be used with any published metabolic model that includes GPR rules. Further, the method can be used with either microarray or RNA-seq data. We demonstrate that a cardiomyocyte-specific model, *iCardio*, with the TIDEs pipeline was able to identify decreased NO synthesis and decreased Neu5Ac synthesis across different heart failure datasets that were not identified using conventional gene-centric approaches, such as gene set enrichment analysis.

## Acknowledgements

The opinions and assertions contained herein are the private views of the authors and are not to be construed as official or as reflecting the views of the U.S. Army or of the U.S. Department of Defense, or The Henry M. Jackson Foundation for Advancement of Military Medicine, Inc. This paper has been approved for public release with unlimited distribution. The authors were supported by the U.S. Army Medical Research and Development Command, Ft. Detrick, MD, as part of the U.S. Army’s Network Science Initiative. Support for this project was provided by the United States Department of Defense (W81XWH-14-C-0054 to JP) and the National Science Foundation Graduate Research Fellowship Program (awarded to BD).

## Author contributions

BD, AW, and JP conceived the study. BD performed the computational modeling and data analysis. BD wrote the initial draft of the manuscript. BD, KR, GK, KV, AW, and JP edited and wrote the final manuscript.

## Data and Code Availability

The code generated in this study is available at https://github.com/csbl/iCardio.

## Methods

### Developing cardiomyocyte-relevant metabolic tasks

Metabolic tasks describe metabolic functions that a tissue or organism is known to be able to catalyze. These metabolic tasks are represented as mathematical constraints on input and output metabolites to the model, where a task is considered to “pass” if there is a feasible flux distribution through the model with the specified constraints. Metabolic tasks have been published with metabolic network reconstructions to demonstrate general metabolic function (Blais et al., 2017; Thiele et al., 2013). For example, *iHsa* was published with 327 metabolic tasks which describe both general metabolism, *i.e*. the generation of ATP from glucose, as well as liver specific metabolism, *i.e*. bile acid synthesis. To expand upon the general metabolic functions in the *iHsa* task list, we curated 94 new, cardiomyocyte-relevant tasks, including 25 that quantitatively describe ATP generation from various carbon sources. Testing these new cardiomyocyte-relevant tasks with *iHsa* resulted in five changes to the network reconstruction (Supplementary Table 1), generating an updated human model which served as the starting point for building the cardiomyocyte-specific model, *iCardio*. Two notable changes were (a) the addition of reactive oxygen species (ROS) formation as 0.1% of the flux through Complex I of the electron transport chain (ETC) as has been done with another model (Smith and Robinson, 2011) and (b) the change from 4 protons to 2.7 protons to generate one ATP molecule to be consistent with recently published data (Watt et al., 2010). The cardiomyocyte-relevant and general metabolic tasks together represent 421 metabolic tasks (Supplemental File 1) that cover a wide range of metabolic functions that both do and do not occur in cardiomyocytes and therefore serve as a resource for curation of the draft *iCardio* model to ensure model functionality (Figure 1).

### Generating and curating a cardiomyocyte-specific metabolic model

The general human model, *iHsa*, was able to pass all metabolic tasks successfully, but achieved a cardiomyocyte-specific task accuracy of 78% since *iHsa* was able to pass the metabolic tasks known to not occur in cardiomyocytes. Since *iHsa* covers all human metabolism, it was necessary to prune reactions from *iHsa* that do not have evidence for presence in cardiomyocytes. To do this, we integrated protein data from the HPA (data available from v18.proteinatlas.org; Uhlén et al., 2015) with *iHsa* (Blais et al., 2017). Various algorithms have been published to generate tissue-specific models from tissue transcriptomics or proteomics data. Here, we implemented 5 of these algorithms (Becker and Palsson, 2008; Jerby et al., 2010; Schultz and Qutub, 2016; Vlassis et al., 2014; Zur et al., 2010) to generate draft cardiomyocyte-specific models. For GIMME, ATP production was used as the objective function. Data was integrated from the HPA, which contains tissue-specific protein expression where each protein is assigned either a high, medium, low, or no expression based on data from antibody-based immunohistochemistry (Uhlén et al., 2015). Code for implementing each algorithm is available at https://github.com/csbl/iCardio. The CORDA algorithm was chosen from among the algorithms given its accuracy for the pre-defined list of cardiomyocyte-specific metabolic tasks (Supplemental File 1, Supplemental Table 2).

The CORDA algorithm takes as an input user-defined high, medium, and negative confidence reactions to produce a model that is (1) consistent (*i.e*. all reactions can carry flux) and (2) maximizes high and medium confidence reactions while minimizing the number of negative confidence reactions. Proteins that corresponded with high, medium, or low/no expression in the heart as indicated in the HPA were included as high (n = 1005), medium (n = 2168), or no confidence reactions (n = 5163) respectively based on the model’s GPR rules. Reactions without GPR rules (~2300 reactions) or reactions associated with no data were included in the negative confidence reactions.

### *Validating* iCardio *using reaction coverage and ATP yields*

*iCardio* was validated qualitatively by determining the number of reactions covered by each metabolic task and quantitatively by comparing ATP yields for common carbon sources between *iCardio* and another metabolic model, *MitoCore* (Smith et al., 2017). Parsimonious flux balance analysis (pFBA) (Lewis et al., 2010) determines the lowest sum of fluxes, and therefore reactions, necessary to complete an objective. Here, pFBA was used to identify the reactions necessary for each metabolic task. Previous work has shown that, in most cases, pFBA produces more physiologically relevant flux distributions compared to flux balance analysis (FBA) or flux-based algorithms which incorporate data (Machado and Herrgård, 2014). As a final, quantitative validation step, we calculated ATP yields for a variety of carbon sources *in silico* as the maximum flux through the ATP synthase reaction for one unit of each carbon source and compared the results to a smaller cardiomyocyte-specific model of mitochondrial metabolism (Smith et al., 2017).

### Analyzing transcriptomics data from heart failure patients

Microarray data (Supplemental Table 3) (Hannenhalli et al., 2006; Kittleson et al., 2005; Liu et al., 2015) from patients undergoing heart transplants for advanced heart failure were downloaded from the Gene Expression Omnibus (GEO) database. Datasets were selected that (a) contained samples for both ischemic and idiopathic heart failure and (b) resulted in at least 50 differentially expressed genes for each type of heart failure. Since all the datasets had been background corrected using RMA, the limma package in R (Ritchie et al., 2015) was used to determine genes that were differentially expressed between healthy hearts and ischemic or idiopathic hearts. Genes corresponding to expression values with an FDR < 0.1 were considered to be differentially expressed and their corresponding fold change was used in subsequent analysis.

### *Identifying <u>T</u>asks <u>I</u>nferred from <u>D</u>ifferential <u>E</u>xpression (TIDEs) using* iCardio *with expression data*

Metabolic tasks and their associated reactions, as identified using *iCardio* with pFBA, were used to identify metabolic functions that are significantly associated with differentially expressed genes in a condition of interest. This method is referred to as <u>T</u>asks <u>I</u>nferred from <u>D</u>ifferential <u>E</u>xpression (TIDEs) (Figure 3). A total of 307 metabolic tasks was used for this analysis, representing the tasks functionally present in *iCardio* from the original task list (Supplemental File 1) that also contained at least one reaction with an associated GPR rule (Supplemental File 3). Reactions that carry flux for each task are identified using a pFBA assumption without previous knowledge of the gene expression data (Figure 3a), as has been done with a related approach (Jerby and Ruppin, 2012). Gene expression log fold changes are overlaid onto reactions in the network using the GPR rules to give each reaction a weight. GPR rules represent the proteins necessary to catalyze a specific reaction through AND or OR relationships. The AND relationships represent a protein complex where different genes encode unique protein subunits necessary for enzyme function while OR relationships represent isozymes. Reactions with complex GPR rules are assigned the maximum absolute fold change for OR relationships and the minimum fold change for AND relationships. For OR relationships where there is a disagreement in the direction of change, *i.e*. where there are genes associated with both a positive and negative fold change, the positive fold change is taken as the reaction weight. The assigned reaction weight values across all reactions in a task are averaged to calculate the task score (Figure 3b). To assign statistical significance to these task scores, the gene expression fold changes are randomized 1000 times amongst the genes measured in each dataset and task scores are recalculated based on the randomized data to create a distribution of task scores. The p-value for each task score that corresponded to the original data is calculated as the number of random task scores greater/less than the original data, depending on how the task score falls relative to the mean randomized task score (Figure 3c). TIDEs are identified as tasks with a p-value < 0.025. A p-value of 0.025 was chosen over a p-value of 0.05 based on the use of a one-sided t-test for calculating significance. Finally, *iCardio* was also used to identify reactions that belonged to each metabolic subsystem in the model. These sets of reactions were also used with the TIDEs method to identify reaction subsystems that were significantly associated with changes in gene expression data. Code is available for re-producing this analysis in MATLAB https://github.com/csbl/iCardio.

### Gene set enrichment analysis

The same gene expression datasets (GSE1869, GSE5406, GSE57345) were also analyzed using gene set enrichment analysis (GSEA) (Subramanian et al., 2005) using the pre-defined gene sets by KEGG pathways. To more closely replicate the TIDE analysis, genes were shuffled within a sample rather than shuffling across samples within each dataset. Pathway enrichment scores with a nominal p-value < 0.05 were defined as statistically significant.

## Supplemental Tables, Figures, and Data

**Supplemental Table 1.**
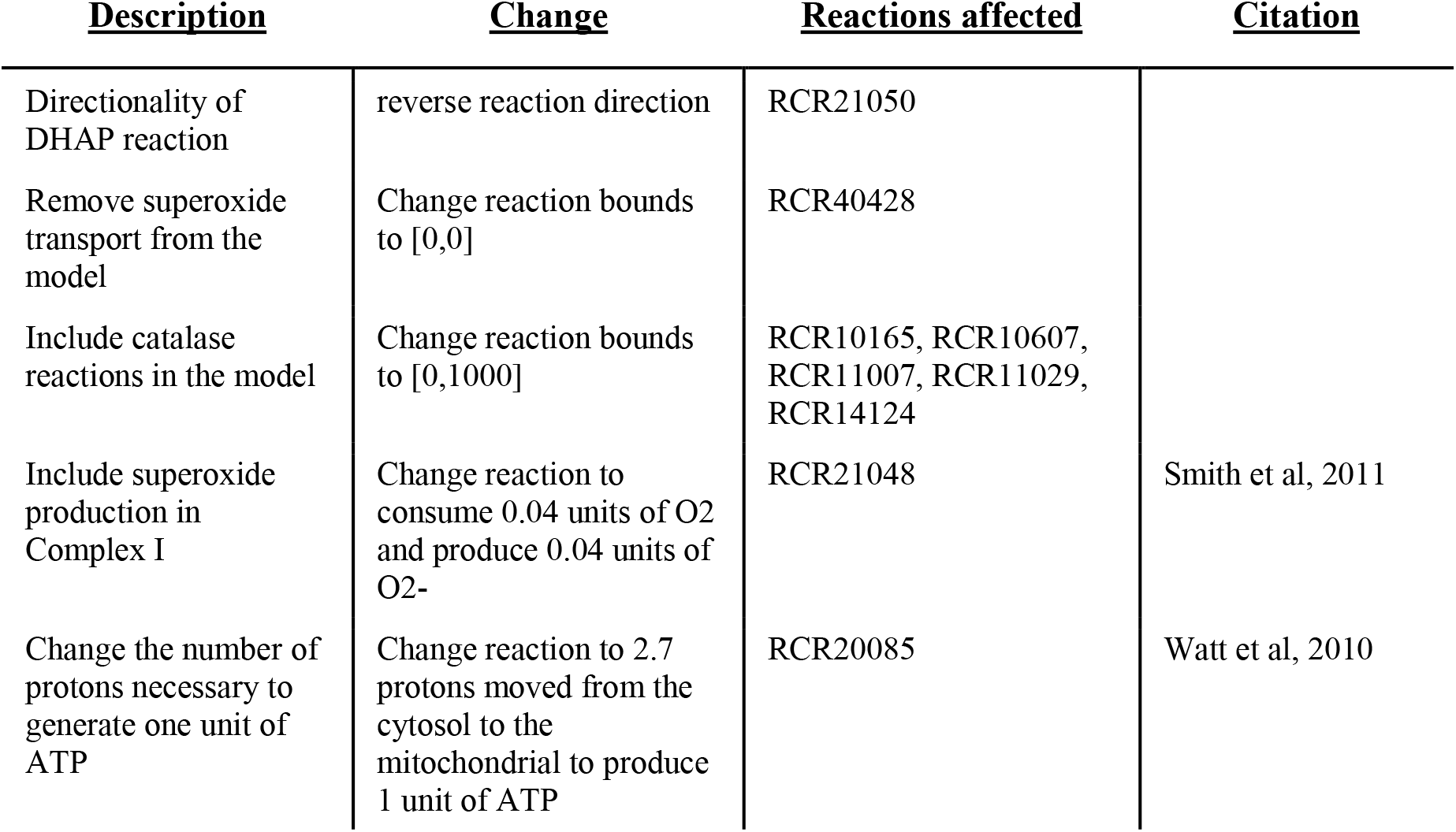
Changes to *iHsa* network reconstruction resulting from metabolic task curation.

**Supplemental Table 2.** Evaluating integration algorithms for construction of draft *iCardio* models. Five different algorithms were used to integrate data from the Human Protein Atlas (HPA) with *iHsa* to produce draft *iCardio* models. Unless otherwise indicated, high and medium proteins, as indicated in the HPA, were used as an input to each algorithm. Cardiomyocyte-specific accuracy was evaluted based on completion of a list of metabolic tasks that covered both general and cardiomyocyte-relevant metabolism (Supplemental File 1).

|  |  | Reactions | Metabolites | Cardiomyocyte-specific task accuracy (%) |
| --- | --- | --- | --- | --- |
| **Original model** | iHsa | 8336 | 5627 | - |
| **Integration algorithms** | **CORDA** | **4133** | **2846** | **89%** |
|  | iMAT | 4434 | 3074 | 82% |
|  | FastCore (high, medium, low) | 4624 | 3215 | 55% |
|  | GIMME | 5526 | 5013 | 37% |
|  | MBA (high, medium, low) | 3709 | 2630 | 37% |
|  | FastCore (only high) | 2671 | 2187 | 36% |
|  | MBA (only high) | 2510 | 2119 | 34% |

**Supplemental Table 3.** Summary of results for individual microarray studies.

| dataset | type of HF | # diseased | # controls | # DEGs | # model DEGs | total genes measured |
| --- | --- | --- | --- | --- | --- | --- |
| GSE1869 | Ischemic | 3 | 6 | 2162 | 278 | 8426 |
| GSE1869 | Idiopathic | 13 | 6 | 6974 | 896 | 8426 |
| GSE57345 | Ischemic | 95 | 136 | 5938 | 808 | 8839 |
| GSE57345 | Idiopathic | 82 | 136 | 5879 | 778 | 8839 |
| GSE5406 | Ischemic | 86 | 16 | 2678 | 392 | 8242 |
| GSE5406 | Idiopathic | 108 | 16 | 2571 | 398 | 8242 |

Supplemental Figure 1. GSEA analysis for all KEGG pathways. Results from the GSEA analysis based for all KEGG pathways, including pathways that were not metabolic. Pathways with no calculated change in any dataset are not shown. As with the TIDEs and KEGG metabolic pathway analysis, results cluster within each dataset rather than by type of heart failure.

Supplemental Figure 2. Subsystem-level analysis using *iCardio* with gene expression data. Using the reaction category assignments already in *iCardio*, reactions associated with each subsystem were determined and processed with the TIDEs pipeline. Subsystems that were significantly increased or decreased based on the gene expression data are shown here.

Supplemental File 1. Metabolic tasks used to evaluate draft *iCardio* models. Includes previously developed *iHsa* tasks and *iCardio* tasks indicating if the task should pass or fail.

Supplemental File 2. Reactions added and removed from the model during manual curation.

Supplemental File 3. Metabolic tasks used for TIDE analysis with results for each GSE study.

## References

Agren, R., Mardinoglu, A., Asplund, A., Kampf, C., Uhlen, M., and Nielsen, J. (2014). Identification of anticancer drugs for hepatocellular carcinoma through personalized genome-scale metabolic modeling. Mol. Syst. Biol. 10.

Bauckneht, M., Ferrarazzo, G., Fiz, F., Morbelli, S., Sarocchi, M., Pastorino, F., Ghidella, A., Pomposelli, E., Miglino, M., Ameri, P., et al. (2017). Doxorubicin Effect on Myocardial Metabolism as a Prerequisite for Subsequent Development of Cardiac Toxicity: A Translational 18F-FDG PET/CT Observation. J. Nucl. Med. 58, 1638–1645.

Becker, S.A., and Palsson, B.O. (2008). Context-specific metabolic networks are consistent with experiments. PLoS Comput. Biol. 4, e1000082.

Blais, E.M., Rawls, K.D., Dougherty, B.V., Li, Z.I., Kolling, G.L., Ye, P., Wallqvist, A., and Papin, J.A. (2017). Reconciled rat and human metabolic networks for comparative toxicogenomics and biomarker predictions. Nat. Commun. 8, 14250.

Blazier, A.S., and Papin, J.A. (2012). Integration of expression data in genome-scale metabolic network reconstructions. Front. Physiol. 3, 299.

Borde, C., Kand, P., and Basu, S. (2012). Enhanced myocardial fluorodeoxyglucose uptake following Adriamycin-based therapy: Evidence of early chemotherapeutic cardiotoxicity? World J. Radiol. 4, 220–223.

Brunk, E., Sahoo, S., Zielinski, D.C., Altunkaya, A., Dräger, A., Mih, N., Gatto, F., Nilsson, A., Preciat Gonzalez, G.A., Aurich, M.K., et al. (2018). Recon3D enables a three-dimensional view of gene variation in human metabolism. Nat. Biotechnol. 36, 272–281.

Doenst, T., Nguyen, T.D., and Abel, E.D. (2013). Cardiac Metabolism in Heart Failure - Implications beyond ATP production. Circ. Res. 113, 709–724.

Duarte, N.C., Becker, S.A., Jamshidi, N., Thiele, I., Mo, M.L., Vo, T.D., Srivas, R., and Palsson, B.Ø. (2007). Global reconstruction of the human metabolic network based on genomic and bibliomic data. Proc. Natl. Acad. Sci. 104, 1777–1782.

Folger, O., Jerby, L., Frezza, C., Gottlieb, E., Ruppin, E., and Shlomi, T. (2011). Predicting selective drug targets in cancer through metabolic networks. Mol. Syst. Biol. 7, 501–501.

Hannenhalli, S., Putt, M.E., Gilmore, J.M., Wang, J., Parmacek, M.S., Epstein, J.A., Morrisey, E.E., Margulies, K.B., and Cappola, T.P. (2006). Transcriptional genomics associates FOX transcription factors with human heart failure. Circulation 114, 1269–1276.

Hayashi, H., Hess, D.T., Zhang, R., Sugi, K., Gao, H., Tan, B.L., Bowles, D.E., Milano, C.A., Jain, M.K., Koch, W.J., et al. (2018). S-Nitrosylation of β-Arrestins Biases Receptor Signaling and Confers Ligand Independence. Mol. Cell 70, 473–487.e6.

Ingwall, J.S. (2004). Transgenesis and cardiac energetics: new insights into cardiac metabolism. J. Mol. Cell. Cardiol. 37, 613–623.

Janardhan, A., Chen, J., and Crawford, P.A. (2011). Altered Systemic Ketone Body Metabolism in Advanced Heart Failure. Tex. Heart Inst. J. 38, 533–538.

Jerby, L., and Ruppin, E. (2012). Predicting Drug Targets and Biomarkers of Cancer via Genome-Scale Metabolic Modeling. Clin. Cancer Res. 18, 5572–5584.

Jerby, L., Shlomi, T., and Ruppin, E. (2010). Computational reconstruction of tissue-specific metabolic models: application to human liver metabolism. Mol. Syst. Biol. 6, 401.

Karlstädt, A., Fliegner, D., Kararigas, G., Ruderisch, H.S., Regitz-Zagrosek, V., and Holzhütter, H.-G. (2012). CardioNet: A human metabolic network suited for the study of cardiomyocyte metabolism. BMC Syst. Biol. 6, 114.

Kittleson, M.M., Minhas, K.M., Irizarry, R.A., Ye, S.Q., Edness, G., Breton, E., Conte, J.V., Tomaselli, G., Garcia, J.G.N., and Hare, J.M. (2005). Gene expression analysis of ischemic and nonischemic cardiomyopathy: shared and distinct genes in the development of heart failure. Physiol. Genomics 21, 299–307.

Kundu, B.K., Zhong, M., Sen, S., Davogustto, G., Keller, S.R., and Taegtmeyer, H. (2015). Remodeling of Glucose Metabolism Precedes Pressure Overload-Induced Left Ventricular Hypertrophy: Review of a Hypothesis. Cardiology 130, 211–220.

Lewis, N.E., Hixson, K.K., Conrad, T.M., Lerman, J.A., Charusanti, P., Polpitiya, A.D., Adkins, J.N., Schramm, G., Purvine, S.O., Lopez-Ferrer, D., et al. (2010). Omic data from evolved E. coli are consistent with computed optimal growth from genome-scale models. Mol. Syst. Biol. 6, 390.

Li, M., Parker, B.L., Pearson, E., Hunter, B., Cao, J., Koay, Y.C., Guneratne, O., James, D.E., Yang, J., Lal, S., et al. (2020). Core functional nodes and sex-specific pathways in human ischaemic and dilated cardiomyopathy. Nat. Commun. 11, 2843.

Li, R., He, H., Fang, S., Hua, Y., Yang, X., Yuan, Y., Liang, S., Liu, P., Tian, Y., Xu, F., et al. (2019). Time series characteristics of serum branched-chain amino acids for early diagnosis of chronic heart failure. J. Proteome Res.

Liu, H., Lv, L., and Yang, K. (2015). Chemotherapy targeting cancer stem cells. Am. J. Cancer Res. 5, 880–893.

Lopaschuk, G.D. (2017). Metabolic Modulators in Heart Disease: Past, Present, and Future. Can. J. Cardiol. 33, 838–849.

Ma, H., Sorokin, A., Mazein, A., Selkov, A., Selkov, E., Demin, O., and Goryanin, I. (2007). The Edinburgh human metabolic network reconstruction and its functional analysis. Mol. Syst. Biol. 3, 135.

Machado, D., and Herrgård, M. (2014). Systematic Evaluation of Methods for Integration of Transcriptomic Data into Constraint-Based Models of Metabolism. PLOS Comput Biol 10, e1003580.

Mardinoglu, A., Gatto, F., and Nielsen, J. Genome-scale modeling of human metabolism – a systems biology approach. Biotechnol. J. 8, 985–996.

Massion, P.B., Feron, O., Dessy, C., and Balligand, J.-L. (2003). Nitric oxide and cardiac function: ten years after, and continuing. Circ. Res. 93, 388–398.

Neubauer, S. (2007). The Failing Heart — An Engine Out of Fuel. N. Engl. J. Med. 356, 1140–1151.

Rawls, K., Dougherty, B.V., and Papin, J. (2020). Metabolic Network Reconstructions to Predict Drug Targets and Off-Target Effects. Methods Mol. Biol. Clifton NJ 2088, 315–330.

Richelle, A., Chiang, A.W.T., Kuo, C.-C., and Lewis, N.E. (2019). Increasing consensus of context-specific metabolic models by integrating data-inferred cell functions. PLoS Comput. Biol. 15, e1006867.

Ritchie, M.E., Phipson, B., Wu, D., Hu, Y., Law, C.W., Shi, W., and Smyth, G.K. (2015). limma powers differential expression analyses for RNA-sequencing and microarray studies. Nucleic Acids Res. 43, e47–e47.

Robaina Estévez, S., and Nikoloski, Z. (2014). Generalized framework for context-specific metabolic model extraction methods. Front. Plant Sci. 5.

Schultz, A., and Qutub, A.A. (2016). Reconstruction of Tissue-Specific Metabolic Networks Using CORDA. PLOS Comput. Biol. 12, e1004808.

Shaked, I., Oberhardt, M.A., Atias, N., Sharan, R., and Ruppin, E. (2016). Metabolic Network Prediction of Drug Side Effects. Cell Syst. 2, 209–213.

Smith, A.C., and Robinson, A.J. (2011). A metabolic model of the mitochondrion and its use in modelling diseases of the tricarboxylic acid cycle. BMC Syst. Biol. 5, 102.

Smith, A.C., Eyassu, F., Mazat, J.-P., and Robinson, A.J. (2017). MitoCore: a curated constraint-based model for simulating human central metabolism. BMC Syst. Biol. 11.

Subramanian, A., Tamayo, P., Mootha, V.K., Mukherjee, S., Ebert, B.L., Gillette, M.A., Paulovich, A., Pomeroy, S.L., Golub, T.R., Lander, E.S., et al. (2005). Gene set enrichment analysis: a knowledge-based approach for interpreting genome-wide expression profiles. Proc. Natl. Acad. Sci. U. S. A. 102, 15545–15550.

Sun, H., Olson, K.C., Gao, C., Prosdocimo, D.A., Zhou, M., Wang, Z., Jeyaraj, D., Youn, J.-Y., Ren, S., Liu, Y., et al. (2016). Catabolic Defect of Branched-Chain Amino Acids Promotes Heart Failure. Circulation 133, 2038–2049.

Swainston, N., Smallbone, K., Hefzi, H., Dobson, P.D., Brewer, J., Hanscho, M., Zielinski, D.C., Ang, K.S., Gardiner, N.J., Gutierrez, J.M., et al. (2016). Recon 2.2: from reconstruction to model of human metabolism. Metabolomics 12, 109.

Taylor, M., Wallhaus, T.R., Degrado, T.R., Russell, D.C., Stanko, P., Nickles, R.J., and Stone, C.K. (2001). An evaluation of myocardial fatty acid and glucose uptake using PET with [18F]fluoro-6-thia-heptadecanoic acid and [18F]FDG in Patients with Congestive Heart Failure. J. Nucl. Med. Off. Publ. Soc. Nucl. Med. 42, 55–62.

Thiele, I., Swainston, N., Fleming, R.M.T., Hoppe, A., Sahoo, S., Aurich, M.K., Haraldsdottir, H., Mo, M.L., Rolfsson, O., Stobbe, M.D., et al. (2013). A community-driven global reconstruction of human metabolism. Nat. Biotechnol. 31, 419–425.

Uhlén, M., Fagerberg, L., Hallström, B.M., Lindskog, C., Oksvold, P., Mardinoglu, A., Sivertsson, Å., Kampf, C., Sjöstedt, E., Asplund, A., et al. (2015). Tissue-based map of the human proteome. Science 347, 1260419.

Vlassis, N., Pacheco, M.P., and Sauter, T. (2014). Fast Reconstruction of Compact Context-Specific Metabolic Network Models. PLOS Comput. Biol. 10, e1003424.

Watt, I.N., Montgomery, M.G., Runswick, M.J., Leslie, A.G.W., and Walker, J.E. (2010). Bioenergetic cost of making an adenosine triphosphate molecule in animal mitochondria. Proc. Natl. Acad. Sci. 107, 16823–16827.

Wende, A.R., Brahma, M.K., McGinnis, G.R., and Young, M.E. (2017). Metabolic Origins of Heart Failure. JACC Basic Transl. Sci. 2, 297–310.

Zhang, A.-D., Dai, S.-X., and Huang, J.-F. (2013). Reconstruction and Analysis of Human Kidney-Specific Metabolic Network Based on Omics Data.

Zhang, C., Chen, J., Liu, Y., and Xu, D. (2019). Sialic acid metabolism as a potential therapeutic target of atherosclerosis. Lipids Health Dis. 18, 173.

Zhao, Y., and Huang, J. (2011). Reconstruction and analysis of human heart-specific metabolic network based on transcriptome and proteome data. Biochem. Biophys. Res. Commun. 415, 450–454.

Zielinski, D.C., Filipp, F.V., Bordbar, A., Jensen, K., Smith, J.W., Herrgard, M.J., Mo, M.L., and Palsson, B.O. (2015). Pharmacogenomic and clinical data link non-pharmacokinetic metabolic dysregulation to drug side effect pathogenesis. Nat. Commun. 6, 7101.

Zur, H., Ruppin, E., and Shlomi, T. (2010). iMAT: an integrative metabolic analysis tool. Bioinformatics 26, 3140–3142.

